# Pinealectomy increases thermogenesis and decreases lipogenesis

**DOI:** 10.1101/813261

**Authors:** Mikyung Kim, So Min Lee, Jeeyoun Jung, Yun Jin Kim, Kyo Chul Moon, Ji Hae Seo, Eunyoung Ha

**Affiliations:** Department of Biochemistry, School of Medicine, Keimyung University, Daegu, Republic of Korea; Clinical Medicine Division, Korea Institute of Oriental Medicine, Daejeon, Republic of Korea

**Keywords:** pinealectomy, melatonin, brown adipose tissue, thermogenesis, lipogenesis

## Abstract

This study was designed to determine the effects of pineal gland-derived melatonin on obesity by employing rat pinealectomy (Pnx) model. After 10 weeks of high-fat diet (HFD) feeding, rats received sham or Pnx surgery followed by 10 weeks normal chow diet (NCD) feeding. Pnx decreased expressions of melatonin receptors, MTNR1A and MTNR1B, in brown (BAT) and white adipose tissues (WAT). Pnx rats showed increased insulin sensitivity compared with those that received sham surgery. Leptin levels were significantly decreased in the serum of Pnx group. In addition, Pnx stimulated thermogenic genes in BAT whereas attenuated lipogenic genes in WAT and the liver. Histologic analyses revealed marked decreased in the size of lipid droplets and increased expressions of UCP1 in BAT and attenuated lipid droplets in the sized and the number in the liver of Pnx group. In conclusion, these results in the current study suggest that Pnx increases thermogenesis in BAT and decreases lipogenesis in WAT and the liver.

## Introduction

Overweight and obesity have been everyday issues for the past decades and statistics proves that they are on a continuous rise worldwide [1]. Obesity is a metabolic syndrome that is associated with a cluster of disorders, ranging from type 2 diabetes mellitus, hypertension and fatty liver disorder [2–4]. Despite the increasing life expectancy in the United States, babies born at the beginning of the twenty-first century are predicted to be the first generation that may have shorter life expectancies than their parents [5]. Many of the risk factors for the overweight and obesity are well defined, but the underlying pathogenesis is not well understood. At present, therapy is aimed at modifying the risk factors, but there are no sustainable therapies for the prevention or even treatment of obesity.

Melatonin is a pineal gland-synthesized neurohormone [6]. It exerts its functions via membrane and nuclear receptors as well as receptor-independent actions. Receptor-mediated neuroendocrine functions of melatonin include the circadian rhythm, sleep, the stress response, the process of aging, and immunity [7,8]. Antioxidant effect of melatonin is a well-established receptor-independent function [9].

Of various functions of melatonin, melatonin promotes energy homeostasis regulating energy balance [10]. Evidences show that melatonin is involved in the regulation of food intake, energy storage, and energy expenditure [11,12]. Melatonin treatment in drinking water or liquid diet reduces body weight and abdominal fat in rats independently of food intake reduction [13–16]. In the zebrafish, melatonin inhibits appetite and stimulates satiety signals in the central nervous system [17]. Evidence also indicates synchronizing function of melatonin with the metabolism in white adipocytes [18].

Although studies with exogenous administration of melatonin show “anti-obesity” effect of melatonin, as referenced above, effects of endogenous melatonin on obesity are controversial and have not been established. Thus, the aim of the current study was to determine the effects of endogenous melatonin, especially pineal gland-derived, on obesity by employing rat pinealectomy model.

## Materials and methods

### Animals

Three-week-old male Wistar rats were purchased from the Laboratory Animal Center of SLC (Japan). The rats were housed in individual cages under controlled temperature (24 ± 1°C), humidity, and lighting (12 : 12 hr light-dark cycle with light at 6:00 a.m.). After one week of acclimatization, rats were fed for 10 weeks high-fat diet (HFD) containing 45% kcal as fat and water *ad libitum* before operation. Rats were randomized to two groups, the sham group with 13 rats and Pnx group with 10 rats. After operation, rats were fed for 10 weeks normal chow diet (NCD) and water *ad libitum*. This study was approved by the Institutional Animal Care and Use Committee (IACUC) of Keimyung University, School of Medicine, Daegu, Korea (KM-2014-16). Body weight, food intake, and water intake were measured twice a week. Peritoneal glucose tolerance test (PGTT) were performed before and after operation on overnight-fasted rats. Rats were killed by isoflurane inhalation at 10 weeks after operation. Tissues including BAT and WAT and serum were harvested for further analyses.

### Surgical procedures

Rats were overnight fasted before surgery. Pnx was performed as described previously [19]. Rats were anesthetized with 5% isoflurane (JW Pharmaceutical Corporation, Seoul, Korea) in 30% oxygen and 70% nitrous oxide. In brief, a sagittal opening was made in the scalp followed by exposure of lambda suture. Skull around the lambda suture was drilled and carefully removed. The pineal gland was then removed with a fine forceps. The removed skull was placed back, and the scalp was sutured. The procedure was completed within 30 min.

### Real-time reverse transcription-polymerase chain reaction (real-time RT-PCR)

Total RNA was extracted with Trizol reagent (Invitrogen, Carlsbad, CA, USA) according to the manufacturer’s protocol. For RT-PCR, 5 μg of total RNA were reverse transcribed for 30 min at 37°C in a reaction mixture containing RNA, 40 U RNase inhibitor (Promega, Madison, WI, USA), 0.5 mM deoxynucleotide triphosphate (Promega, Madison, WI, USA),2 μM random hexamer primers, 5 x AMV reverse transcriptase reaction buffer and 30 U AMV reverse transcriptase (Promega, Madison, WI, USA). Real-time RT-PCR analysis was performed using the SYBR Green PCR Master Mix (TOYOBO, Osaka, Japan) and real-time RT-PCR parameters were used as follows: initial denaturation 95°C for 30 s, amplification 45 cycles were performed at 95°C for 5 s, 60°C for 10 s, and 72°C for 15 s, melting curve 95°C for 10 s, 65°C for 1 min, cooling 37°C for 10 min with gene-specific primers. The relative expression of each gene was normalized against beta-actin. The samples were assayed on a Light Cycler 480 (Roche, Basel, Switzerland) instrument and the concentration was calculated as copies per μL using the standard curve. Primer sequences (Macrogen, Seoul, Korea) are listed in Table 1.

**Table 1.**
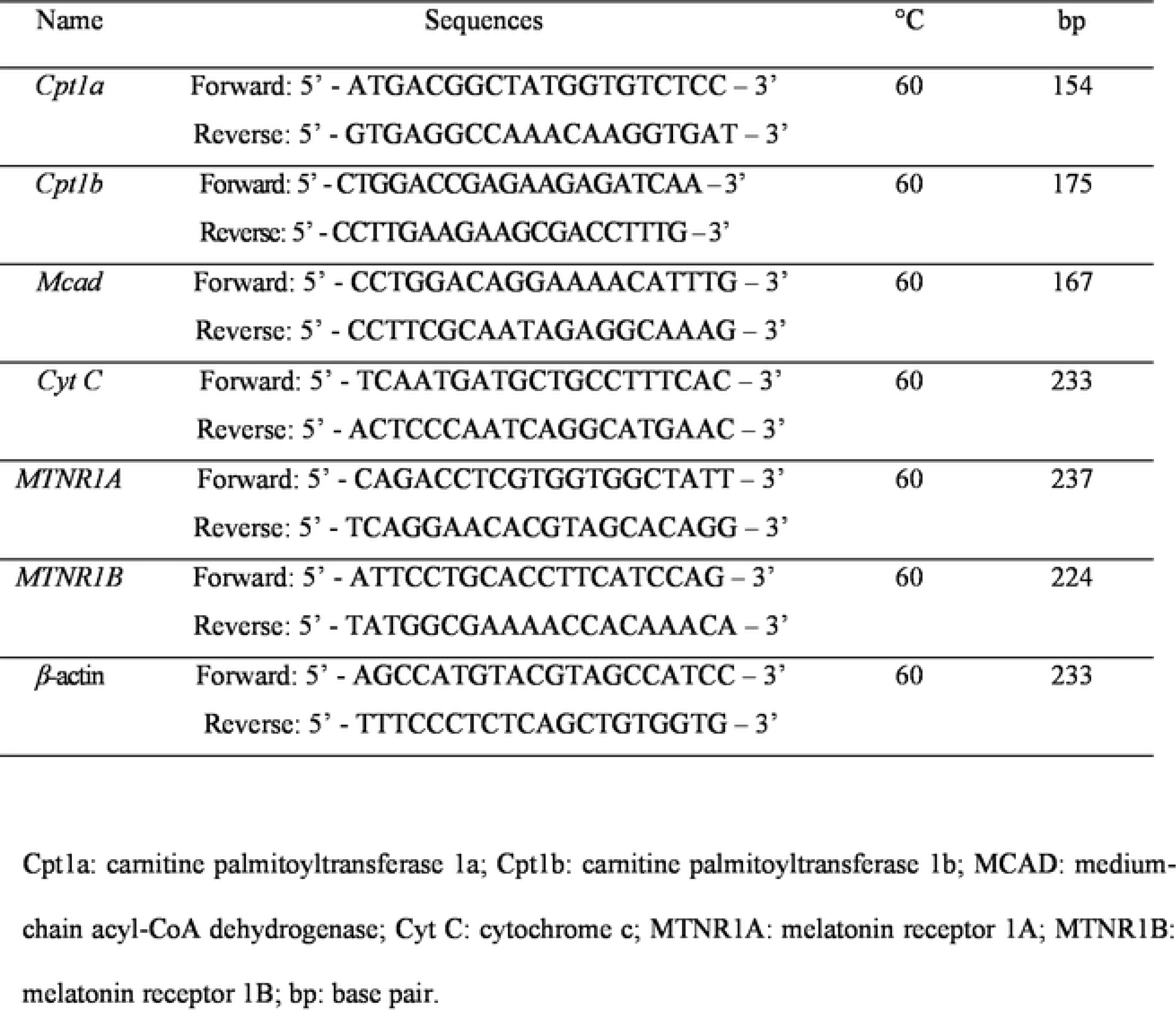
Primer sequences

### Western blot analysis

Harvested tissues were subjected to SDS-PAGE and immunoblotted. Briefly, tissues were lysed in ice-cold lysis buffer [50 mM Tris-HCl (pH 7.4), 25 mM EDTA (pH 8.0), 650 mM NaCl, 5% Triton X-100] containing protease inhibitors (200 mM phenylmethylsulfonyl fluoride, 100 μg/mL leupeptin, 10 μg/mL pepstatin, 1 μg/mL aprotinin, and 2 mM EDTA). The lysates were centrifuged for 20 min at 7,500 x g at 4°C and the supernatant was collected. Proteins were separated by SDS-PAGE and transferred to nitrocellulose membrane (GE Healthcare, Little Chalfont, UK). The membrane was blocked in 5% skim milk in TBS-T (Tris buffered saline plus 0.1% Tween 20) before incubation for overnight at 4°C with primary antibody of anti-UCP1, anti-PGC1α, anti-DIO2 (1: 1,000 dilution, Abcam, Cambridge, USA) and anti-SIRT1 (1: 1,000 dilution, Santa Cruz Biotechnology, Santa Cruz, CA, USA). Beta-actin (1: 1,000 dilution, Sigma, St. Louis, MO, USA) was used as an internal control. Membrane was then washed in TBS-T and incubated with horseradish peroxidase-conjugated secondary antibodies (1: 1,000 dilution, Santa Cruz Biotechnology, Santa Cruz, CA, USA). Protein bands were detected using Super Signal West Pico Chemiluminescent Substrate (Thermo Fisher Scientific, Boston, MA, USA). The protein-specific signals were detected using LAS-3000 (Fujifilm, Tokyo, Japan).

### Histological assessment of BAT

BAT of each rats was fixed with 10% formalin and embedded in paraffin and sectioned. The sections were stained with hematoxylin and eosin (H&E) or immunohistochemistry (IHC) for light microscopic examination. For assessment of UCP1 staining, 5 μm sections were permeabilized in PBS, incubated in 10 mM sodium citrate buffer with pH 6.0 for 100°C, 10 min, and the with anti-UCP1 (1: 3,000 dilution, Abcam, Cambridge, USA) antibody incubation for overnight at 4°C. Sections were then incubated with secondary antibody (1: 200 dilution, Santa Cruz Biotechnology, Santa Cruz, CA, USA) for 1 hr at room temperature and followed by staining with diaminobenzidine chromogen (Vector Laboratories, CA, USA) and counterstaining with hematoxylin. The stained sections were examined under microscopy (x 200) and all histological assessments (Nikon, Tokyo, Japan) were made by a pathologist.

### Enzyme-linked immunosorbent assay (ELISA)

Blood samples were allowed to clot for 2 hr at room temperature before centrifuging for 20 min at 2,000 x g. Leptin levels were measured by ELISA kit (R&D System, Minneapolis, MN, USA) in serum. Enzyme levels in supernatants were measured according to manufacturer’s instructions. Serum samples were then diluted 1: 10 in diluent solution and incubated in the plates at room temperature for 2 hr, after which the plates were washed five times. The detection rat leptin conjugate was incubated for 2 hr at room temperature. The plates were washed five times and substrate solution was incubated for 30 min, after which stop solution was added. Standard serum from rat was added to each plate in serial dilutions, and a standard curve was constructed to assign arbitrary units. The absorbance values were determined with an ELISA microplate reader (Biochrom, Cambridge, UK) measured in wavelengths of 450 nm and then 570 nm for corrections.

### Statistical analysis

Data were analyzed by Students *t*-test for unpaired comparisons. The accepted level of significance was preset as P value < 0.05. Each experiment was executed at least three times in duplicate. All Data are given in terms of relative values and presented as means ± standard deviation (SD).

## Results

### Pnx increased insulin sensitivity

Body weights before Pnx were 465 ± 22.3 g. Body weights at the end of experiment were 542 ± 17.5 g in sham and 529 ± 28.1 g in Pnx group. Although not statistically different, the degree of increase in body weights in Pnx group was consistently less than that in sham group until the end of experiment (Figure 1A). PGTT before the operation revealed impaired insulin responses. PGTT after the operation revealed that glucose levels at 60, 90, and 120 min in Pnx group were lower than those in sham group, indicating an improvement of insulin sensitivity in Pnx group (Figure 1B). Food and water intakes per 100g body weight did not differ between two groups (Figure 1C and D). Since insulin resistance is associated with elevated plasma leptin levels (Segal, et al. 1996), we then investigated circulating leptin levels in sham and Pnx groups (Figure 1E). Circulating leptin levels in Pnx group noticeably decreased (159.79 ± 13.39 pg/mL, p < 0.01) compared with those in sham group (344.67 ± 48.64 pg/mL, p < 0.05), implicating increased insulin sensitivity by Pnx.

**Figure 1.**
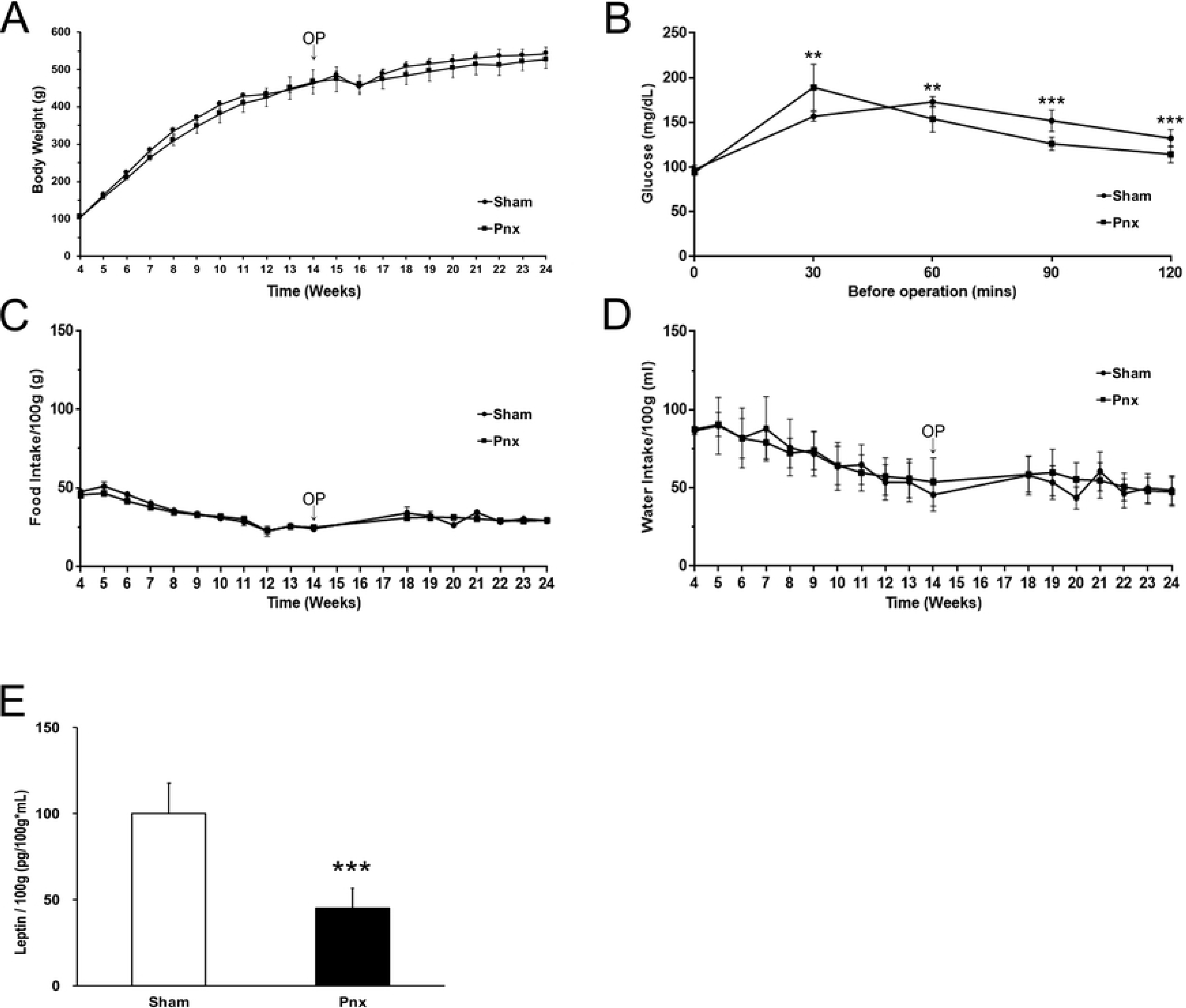
Changes in body weight (A), PGTT (B), food intake (C), water intake (D) and plasma leptin levels (E). Values are mean ± SD. Pnx: pinealectomy; PGTT: peritoneal glucose tolerance test; OP: operation. \*\**P* < 0.01, \*\*\**P* < 0.001, compared with sham group.

### Pnx decreased MTNR1A and MTNR1B expressions in WAT and BAT

We examined the expression levels of melatonin receptors, MTNR1A and MTNR1B in WAT and BAT to determine the effect of Pnx on the expressions of melatonin receptors Figure 2A and B). Expectantly, we found that both mRNA and protein expression levels of MTNR1A and MTNR1B decreased in Pnx group compared with those in sham group in WAT (Figure 2).

**Figure 2.**
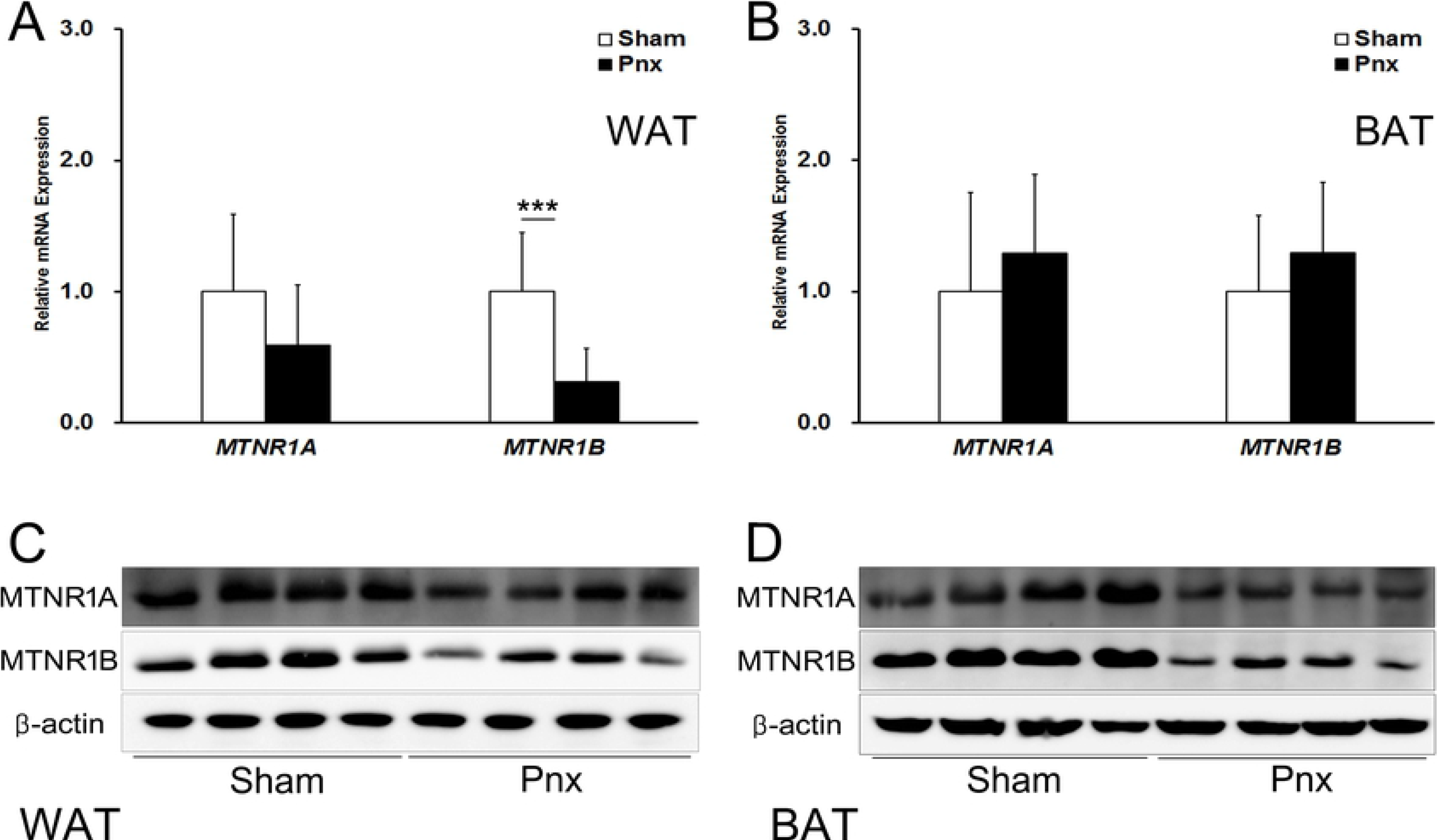
Expressions of melatonin receptors, MTNR1A and MTNR1B, in WAT and BAT. (A) mRNA expressions of MTNR1A and MTNR1B in WAT and (B) BAT were examined by real-time RT-PCR and normalized to β-actin. (C) Protein expressions of MTNR1A and MTNR1B in WAT and (D) BAT were examined by Western blot analysis. Values are mean ± SD. Pnx: pinealectomy; WAT: white adipose tissue; BAT: brown adipose tissue; real-time RT-PCR: real-time reverse transcription-polymerasechain reaction; MTNR1A: melatonin receptor 1A; MTNR1B: melatonin receptor 1B. \**P* < 0.05, \*\*\**P* < 0.001, compared with sham group.

### Pnx stimulated Thermogenesis in BAT

Based on the results that showed increased insulin sensitivity and decreased MTNR1A and MTNR1B expressions in WAT and BAT in Pnx group, we hypothesized that Pnx improved insulin sensitivity via stimulating thermogenesis in BAT and regulating lipogenesis in WAT. To explore our hypothesis, we determined thermogenic genes in BAT (Figure 3A). Figure 3A showed pronounced increases in the expressions (upper panel) and quantifications (lower panel) of thermogenic genes, sirtuin 1 (SIRT1), peroxisome proliferator-activated receptor gamma coactivator-1α (PGC1α), uncoupling protein 1 (UCP1), and iodothyronine deiodinase 2 (DIO2) in Pnx groups compared with those in sham group. Accordingly, we also found increased mRNA expressions of PGC1α, UCP1, and DIO2, PR domain-containing 16 (PRDM16) in Pnx group (Figure 3B and C). We observed, however, no difference in the level of PPARγ, the master regulator of adipogenesis.

**Figure 3.**
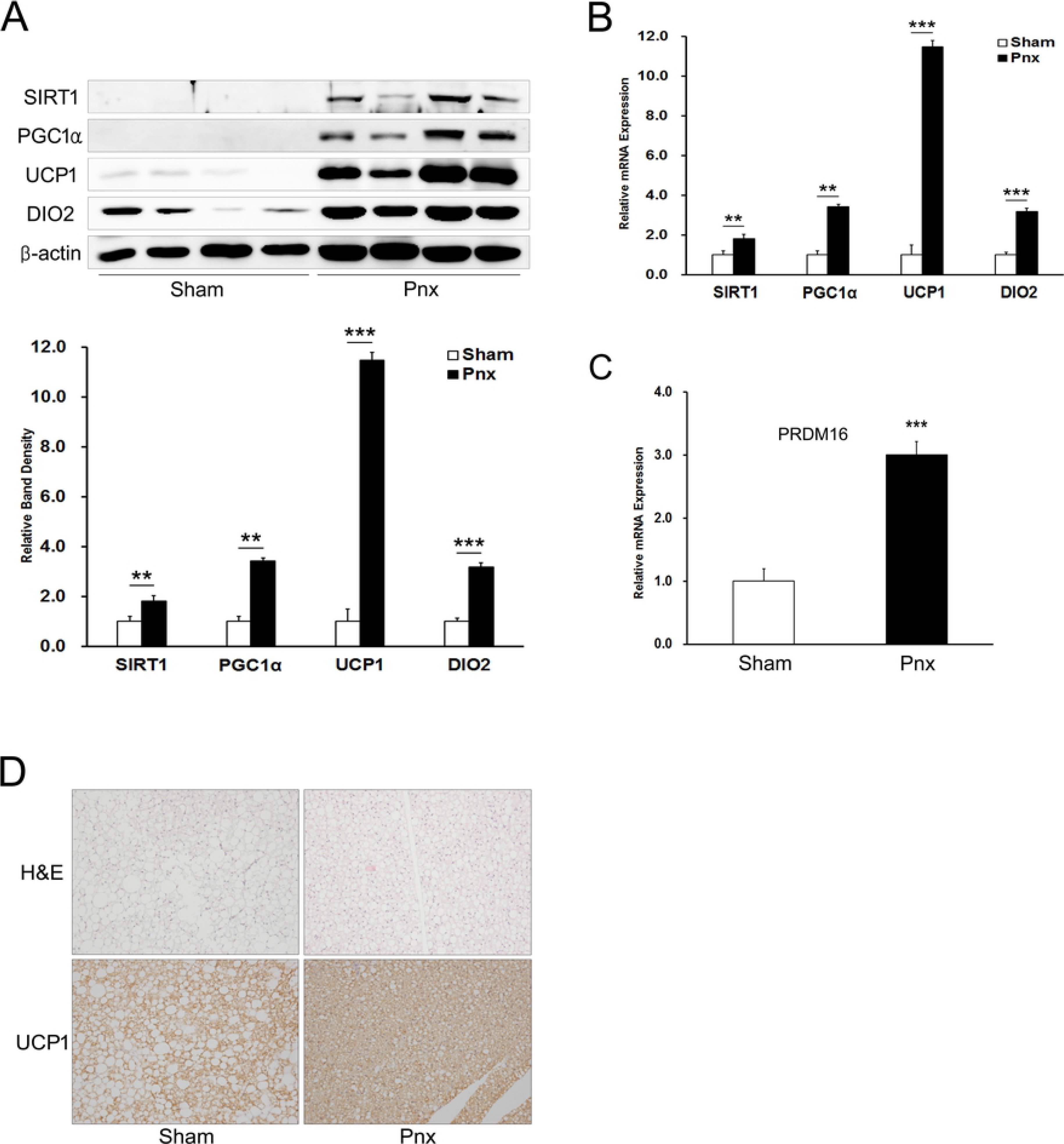
Thermogenic genes and PRDM16 in BAT. (A) Protein expressions of SIRT1, PGC1α, UCP1 and DIO2 were examined by Western blot analysis (upper panel) and relative band density (lower panel). (B) mRNA expressions of PGC1α, UCP1, DIO2, PPARγ, and (C) PRDM16 were examined by real-time RT-PCR and normalized to β-actin. (D) Changes of lipid droplets and UCP1 expressions in BAT. Representative images of BAT from Sham group and Pnx group stained with H&E (upper pannel) and immune-histochemical images stained with UCP1 (lower pannel). Original magnification (x 200). Values are mean ± SD. Pnx: pinealectomy; BAT: brown adipose tissue; real-time RT-PCR: real-time reverse transcription-polymerase chain reaction; PGC1α: peroxisome proliferator-activated receptor gamma coactivator 1-alpha; UCP1: uncoupling protein 1; PPARγ: peroxisome proliferator-activated receptor gamma; DIO2: deiodinase 2; Prdm16: PR domain 16; H&E: hematoxylin and eosin. \**P* < 0.05, \*\*\**P* < 0.001, compared with sham group.

To further validate thermogenic effect of Pnx, we then examined histological changes and UCP1 expressions in BAT (Figure 3D). Histologic analysis of BAT by H & E staining revealed marked decrease in the size of lipid droplets in Pnx group (Figure 3D upper right panel). Consistent with the result in Figure 3A and B, immunohistochemical staining showed pronounced increase in the expression of UCP1in Pnx group compared with sham group (Figure 3D lower right panel).

### Pnx down-regulated lipogenic genes in WAT and the liver

Next, we examined the possible regulatory effects of Pnx in WAT by determining the expression levels of lipogenic genes (Figure 4A). We found that Pnx caused a noteworthy reduction of SREBP1c, the master regulatory transcription factor for lipogenesis. Consistent with the decreased expression of sterol regulatory element binding protein 1c (SREBP1c), we also found in Pnx group decreased expression of fatty acid synthase (FASN), a key lipogenic enzyme in the first step of *de novo* lipogenesis. Moreover, Pnx decreased the expression of stearoyl-CoA desaturase (SCD1), which catalyzes the unsaturation of fatty acids forming a double bond in stearoyl-CoA. These results suggest that Pnx inhibited lipogenic pathway in WAT. We next determined major mitochondrial biogenesis genes PGC1α and cytochrome c (Cyt c). Contrary to the expression in BAT, the expression of PGC1α in WAT was decreased, which might indicate attenuated mitochondrial biogenesis. Decreased expressions of genes involved in fatty acid oxidation (CPT1b and medium-chain acyl-CoA dehydrogenase), may supports attenuated mitochondrial biogenesis (Figure 4B).

**Figure 4.**
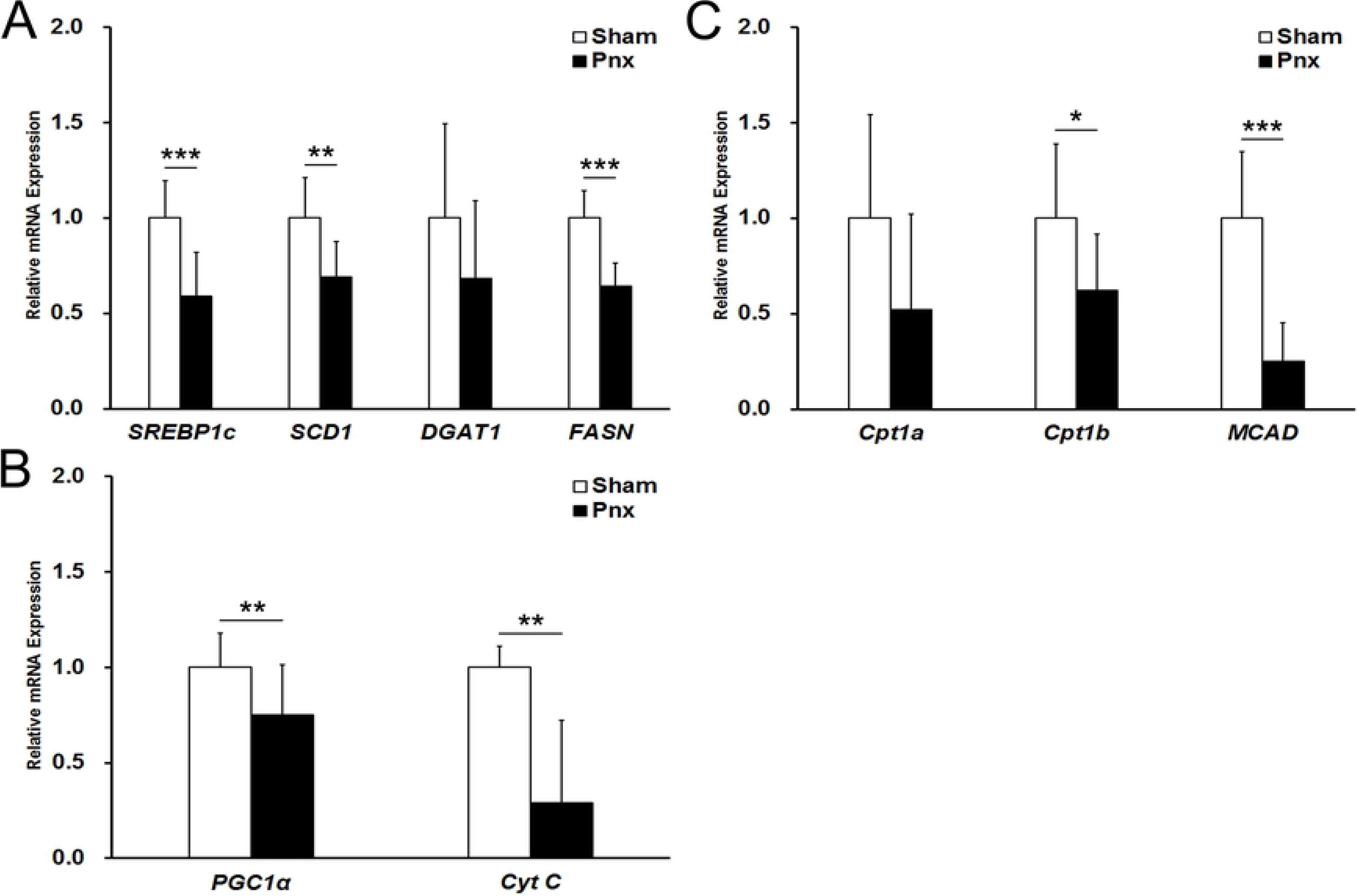
Lipogenesis genes in WAT. (A) mRNA expressions of lipogenic genes, SREBP1c, SCD1, DGAT1 and FASN, in WAT were examined by real-time RT-PCR and normalized to β-actin. (B) mRNA expressions of mitochondrial biogenesis genes, PGC1α and Cyt C, in WAT were examined by real-time RT-PCR and normalized to β-actin. (C) mRNA expressions of fatty acid oxidation genes, CPT1a, CPT1b and MCAD, in WAT were examined by real-time RT-PCR and normalized to β-actin. Values are mean ± SD. Pnx: pinealectomy; WAT: white adipose tissue; real-time RT-PCR: real-time reverse transcription-polymerase chain reaction; SREBP1c: sterol regulatory element-binding protein 1; SCD1: stearoyl-CoA desaturase 1; DGAT1: diacylglycerol O-acyltransferase 1; FASN: fatty acid synthase; CPT1a: carnitine palmitoyltransferase 1a; CPT1b: carnitine palmitoyltransferase 1b; MCAD: medium-chain acyl-CoA dehydrogenase; PGC1α: peroxisome proliferator-activated receptor gamma coactivator 1-alpha; Cyt C: cytochrome c. \**P* < 0.05, \*\**P* < 0.01, \*\*\**P* < 0.001, compared with sham group.

Since liver is the organ mostly responsible for lipogenesis, we further examined the expression levels of lipogenic genes in the liver. Histologic analysis of the liver by H & E staining also showed attenuated lipid droplets both in the size and the number in Pnx group compared with sham group (Figure 5A). In line with the results of the expression levels in WAT, we found that Pnx induced a significant decrease in SREBP1c. Expressions of SREPB1c target genes, SCD1, FASN, and DGAT2, although not statistically significant, showed tendencies to decrease in Pnx group (Figure 5B). Expressions of genes in mitochondrial biogenesis and fatty acid oxidation, except CPT1a, showed no difference between sham and Pnx groups (Figure 5C and D).

**Figure 5.**
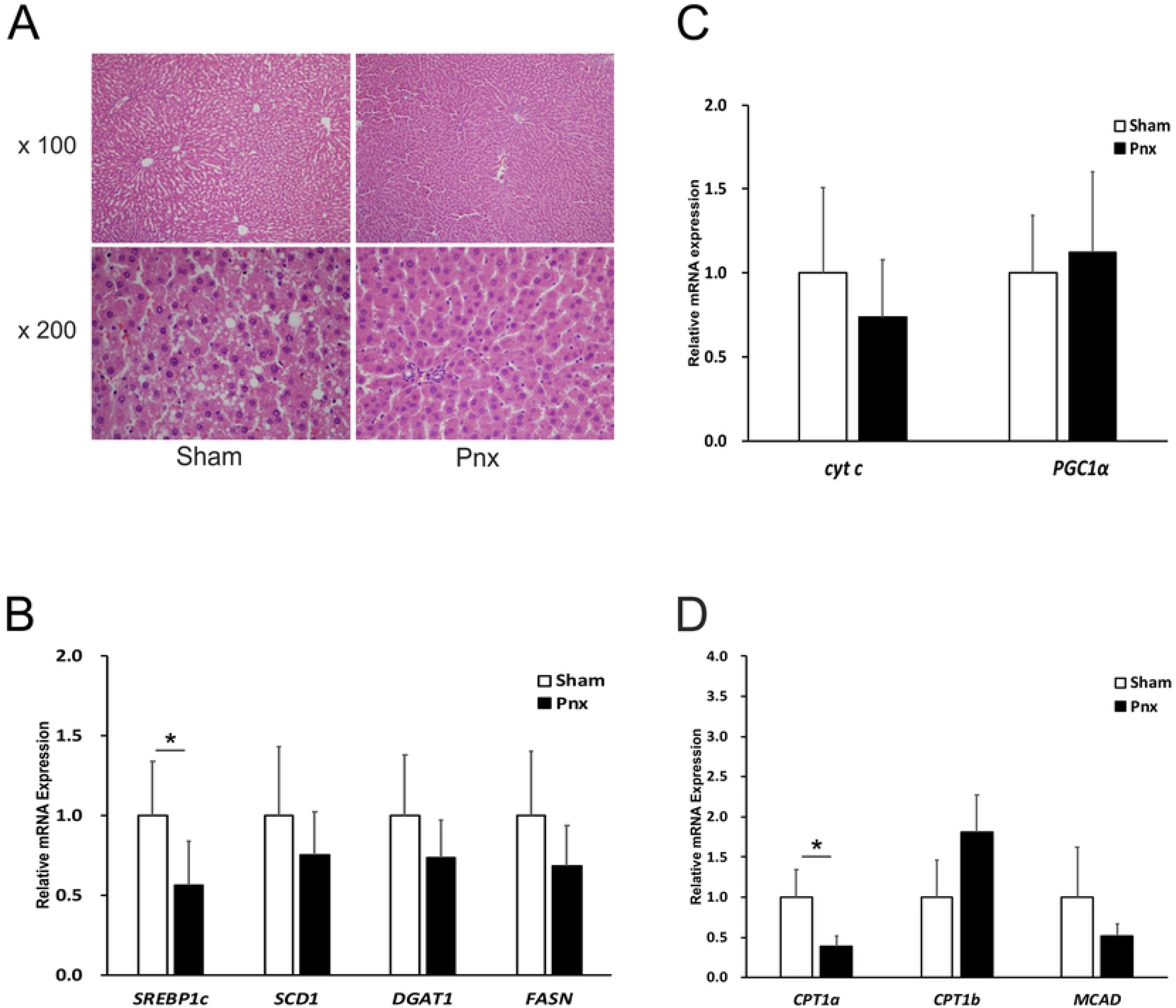
Changes in hepatic histology and lipogenesis genes in the liver. (A) Liver sections were stained with H&E staining. Original magnification (x 100 upper panel and x200 lower panel). (B) mRNA expressions of lipogenic genes, SREBP1c, SCD1, DGAT1 and FASN, in the liver were examined by real-time RT-PCR and normalized to β-actin. (C) mRNA expressions of mitochondrial biogenesis genes, PGC1α and Cyt C, in the liver were examined by real-time RT-PCR and normalized to β-actin. (D) mRNA expressions of fatty acid oxidation genes, CPT1a, CPT1b and MCAD, in the liver were examined by real-time RT-PCR and normalized to β-actin. Values are mean ± SD. Pnx: pinealectomy; real-time RT-PCR: real-time reverse transcription-polymerase chain reaction; SREBP1c: sterol regulatory element-binding protein 1; SCD1: stearoyl-CoA desaturase 1; DGAT1: diacylglycerol O-acyltransferase 1; FASN: fatty acid synthase; Cpt1a: carnitine palmitoyltransferase 1a; Cpt1b: carnitine palmitoyltransferase 1b; MCAD: medium-chain acyl-CoA dehydrogenase; PGC1α: peroxisome proliferator-activated receptor gamma coactivator 1-alpha; Cyt C: cytochrome c; H&E: hematoxylin and eosin. *P < 0.05, **P < 0.01, compared with sham group.

## Discussion

Extensive studies show that treatment of melatonin reduces body weight and food intake [13–16]. In the current study, however, we showed that Pnx stimulated thermogenesis in BAT and attenuated lipogenesis in WAT and the liver of HFD fed rats without any significant changes in body weight compared with control rats.

We observed no difference in body weight and food intake in Pnx group compared with those in control group, which may implicate that endogenously synthesized pineal melatonin does not affect systemic energy balance. To support our result, a very recent study showed no difference in body weight and food intake between normal chow diet-fed control and Pnx rats [20]. The study also indicated that exogenous treatment of melatonin to Pnx group resulted in body weight and food intake decrease, which suggests that exogenous, rather than endogenous, melatonin as a possible contributing factor for the decreases in body weight and food intake.

Intriguingly, despite no difference in body weight changes, we observed that Pnx increased insulin sensitivity evidenced by faster disposal of plasma glucose and lower plasma leptin level in Pnx. These results are in contrast to the previous report that showed Pnx induces insulin resistance [21]. Lima et al. demonstrated that pinealectomy causes glucose intolerance and decreases adipose cell responsiveness to insulin in normal chow diet-fed rats. Since HFD-fed rats were employed in this study, we speculated the possible interaction between diet and the absence of pineal melatonin so as to affect systemic metabolism. Given the well-established facts that leptin levels increase in obesity and are associated with insulin resistance, decreased leptin levels in Pnx group compared with that in control HFD-fed group further validates Pnx induced insulin sensitivity [22–24].

UCP1, also known as thermogenin and is predominantly expressed in BAT, is a mitochondrial protein that regulates dissipation of excess energy via the uncoupling of oxidative phosphorylation. It is also positively associated with insulin sensitivity [25]. In the current study, we showed that Pnx markedly increased the expressions of UCP1 in BAT. To support the increased activity of UCP1 in Pnx, we also found increased expressions of PGC1α, DIO2, and SIRT1 in Pnx group. The activity of UCP1 is involved in several transcript factors and gene, such as PGC1α, DIO2, SIRT1, and PRDM16. PGC1α is a master nuclear transcription factor that controls the expression of the thermogenic genes [26]. There was markedly increased expression of PGC1α, which implicates PGC1α-induced thermogenic genes. DIO2 is another well-known master regulatory molecule that controls the expression of the thermogenic genes [27]. In this study, we also observed that Pnx stimulated the expression of DIO2 compared with that in sham group. With the increased expressions of PGC1α and DIO2, expression of thermogenic gene UCP1 was expectantly increased in BAT of Pnx group. Furthermore, DIO2 activity generates a significant fraction of circulating T3 in rats and then enhances the cAMP-generated acute increase in UCP1 mRNA via increased UCP1 gene transcription [28]. Prdm16 is recently identified regulatory molecule to induce brown adipogenecity [29]. And the expression of Prdm16 in BAT of Pnx was increased which implicates Pnx induced BAT. More intriguingly, we found that the expression of SIRT1 in Pnx increased the expressions of SIRT1 in BAT, which further implicates the Pnx-induced insulin sensitivity might be mediated via BAT activation [30].

## Conclusions

Pnx decreased lipogenesis in WAT and the liver. Here, we report that expressions of lipogenic genes were decreased in WAT and the liver of Pnx group compared with those in WAT of sham group, indicating Pnx attenuates lipogenesis via as yet unidentified mechanism. These results indicate that Pnx may inhibit the lipogenic activity by inhibiting the lipogenic gene. This is supported by decreases in gene expressions of mitochondrial biogenesis genes, PGC1α and Cyt C. In conclusion, Pnx in rats increases thermogenesis in BAT and decreases lipogenesis in WAT, leading to increased insulin sensitivity.

## Author Contributions

**Conceptualization:** Eunyoung Ha, Ji Hae Seo

**Data curation:** Mikyung Kim, So Min Lee

**Formal analysis:** Mikyung Kim, So Min Lee, Jeeyoun Jung, Yun Jin Kim

**Funding acquisition:** Eunyoung Ha, Ji Hae Seo

**Investigation:** Mikyung Kim, So Min Lee, Kyo Chul Moon, Eunyoung Ha

**Methodology:** Mikyung Kim, So Min Lee, Jeeyoun Jung

**Supervision:** Eunyoung Ha, Jeeyoun Jung, Kyo Chul Moon, Ji Hae Seo

**Visualization:** Kyo Chul Moon

**Writing-original draft:** Mikyung Kim, So Min Lee, Kyo Chul Moon

**Writing-review & editing:** Eunyoung Ha, Ji Hae Seo

## References

1. Ng M, Fleming T, Robinson M, Thomson B, et al. Global, regional, and national prevalence of overweight and obesity in children and adults during 1980-2013: a systematic analysis for the Global Burden of Disease Study. Lancet 2014; 384: 766–781.

2. Berrington de Gonzalez A, Hartge P, Cerhan JR, et al. Body-mass index and mortality among 1.46 million white adults. N Engl J Med 2010; 363: 2211–2219.

3. Renehan AG, Tyson M, Egger M, et al. Body-mass index and incidence of cancer: a systematic review and meta-analysis of prospective observational studies. Lancet 2008; 371: 569–578.

4. Whitlock G, Lewington S, Sherliker P, et al. Body-mass index and cause-specific mortality in 900 000 adults: collaborative analyses of 57 prospective studies. Lancet 2009; 373: 1083–1096.

5. Olshansky SJ, Passaro DJ, Hershow RC, et al. A potential decline in life expectancy in the United States in the 21st century. N Engl J Med 2005; 352: 1138–1145.

6. Stehle JH, Saade A, Rawashdeh O, et al. A survey of molecular details in the human pineal gland in the light of phylogeny, structure, function and chronobiological diseases. J Pineal Res 2011; 51: 17–43.

7. Cardinali DP, Golombek DA, Rosenstein RE, et al. Melatonin site and mechanism of action: single or multiple? J Pineal Res 1997; 23: 32–39.

8. Galano A, Tan DX and Reiter RJ. Melatonin as a natural ally against oxidative stress: a physicochemical examination. J Pineal Res 2011; 51: 1–16.

9. Reiter RJ, Tan DX, Osuna C, et al. Actions of melatonin in the reduction of oxidative stress. J Biomed Sci 2000; 7: 444–458.

10. Cipolla-Neto J, Amaral F, Afeche S, et al. Melatonin, energy metabolism, and obesity: a review. J Pineal Res 2014; 56: 371–381.

11. Bartness TJ and Wade GN. Body weight, food intake and energy regulation in exercising and melatonin-treated Siberian hamsters. Physiol Behav 1985; 35: 805–808.

12. Bubenik G and Pang SF. The role of serotonin and melatonin in gastrointestinal physiology: Ontogeny, regulation of food intake, and mutual serotonin-melatonin feedback. J Pineal Res 1994;16: 91–99.

13. Bojkova B, Orendáš P, Friedmanova L, et al. Prolonged melatonin administration in 6-month-old Sprague-Dawley rats: metabolic alterations. Acta Physiol Hung 2008; 95: 65–76.

14. Puchalski SS, Green JN and Rasmussen DD. Melatonin effect on rat body weight regulation in response to high-fat diet at middle age. Endocrine 2003; 21: 163–167.

15. Ríos-Lugo MJ, Jiménez-Ortega V, Cano-Barquilla P, et al. Melatonin counteracts changes in hypothalamic gene expression of signals regulating feeding behavior in high-fat fed rats. Horm Mol Biol Clin Investig 2015; 21: 175–183.

16. Wolden-Hanson T, Mitton D, McCants R, et al. Daily melatonin administration to middle-aged male rats suppresses body weight, intraabdominal adiposity, and plasma leptin and insulin independent of food intake and total body fat. Endocrinology 2000; 141: 487–497.

17. Piccinetti CC, Migliarini B, Olivotto I, et al. Appetite regulation: the central role of melatonin in Danio rerio. Horm Behav 2010; 58: 780–785.

18. Alonso-Vale MIC, Andreotti S, Mukai PY, et al. Melatonin and the circadian entrainment of metabolic and hormonal activities in primary isolated adipocytes. J Pineal Res 2008; 45: 422–429.

19. Maganhin CC, Simões RS, Fuchs LF, et al. Rat pinealectomy: a modified direct visual approach. Acta Cir Bras 2009; 24: 321–324.

20. Buonfiglio D, Parthimos R, Dantas R, et al. Melatonin absence leads to long-term leptin resistance and overweight in rats. Front Endocrinol 2018; 9: 122.

21. Lima FB, Machado UF, Bartol I, et al. Pinealectomy causes glucose intolerance and decreases adipose cell responsiveness to insulin in rats. Am J Physiol 1998; 275: E934–E941.

22. Considine RV, Sinha MK, Heiman ML, et al. Serum immunoreactive-leptin concentrations in normal-weight and obese humans N Engl J Med 1996; 334: 292–295.

23. Handjieva-Darlenska T and Boyadjieva NJ. The effect of high-fat diet on plasma ghrelin and leptin levels in rats. J Physiol Biochem 2009; 65: 157–164.

24. Segal KR, Landt M and Klein S. Relationship between insulin sensitivity and plasma leptin concentration in lean and obese men. Diabetes 1996; 45: 988–991.

25. Chondronikola M, Volpi E, Børsheim E, et al. Brown adipose tissue improves whole body glucose homeostasis and insulin sensitivity in humans. Diabetes 2014; 63: 4089–4099.

26. Seale P, Kajimura S, Yang W, et al. Transcriptional control of brown fat determination by PRDM16. Cell Metab 2007; 6: 38–54.

27. Bartelt A and Heeren J. Adipose tissue browning and metabolic health. Nat Rev endocrinol 2014; 10: 24–36.

28. De Jesus LA, Carvalho SD, Ribeiro MO, et al. The type 2 iodothyronine deiodinase is essential for adaptive thermogenesis in brown adipose tissue. J Clin Invest 2001; 108: 1379–1385.

29. Giralt M and Villarroya F. White, brown, beige/brite: different adipose cells for different functions? Endocrinology 2013; 154: 2992–3000.

30. Boutant M, Joffraud M, Kulkarni SS, et al. SIRT1 enhances glucose tolerance by potentiating brown adipose tissue function. Mol Metab 2015; 4: 118–131.

